# Enhancing Sensitivity in Targeted Single-Cell Proteomics by Coupling a Dual Ion Funnel Interface with Triple Quadrupole Mass Spectrometer

**DOI:** 10.64898/2026.01.02.697431

**Authors:** Sehong Min, Pearl Kwantwi-Barima, Issac K. Attah, Thomas L. Filmore, Matthew J. Gaffrey, Reta Birhanu Kitata, Yehia M. Ibrahim, Tujin Shi

## Abstract

Single-cell proteomics (SCP) has emerged as a powerful approach for understanding cellular heterogeneity and biological processes at unprecedented resolution. However, the extremely limited protein content of individual cells (femtogram to picogram levels) pushes current mass spectrometry instrumentation to its sensitivity limits, creating a critical analytical bottleneck. While selected reaction monitoring (SRM) using triple quadrupole (QqQ) instruments offers advantages in sensitivity and reproducibility for targeted proteomics quantification, SRM still struggles with sensitivity for quantification of moderate- or low-abundance proteins from single-cell sample amounts. Here, we report the development and systematic evaluation of a dual ion funnel interface designed to address the sensitivity limitation by significantly enhancing ion transmission efficiency in commercial QqQ mass spectrometers. The dual ion funnel interface, composed of a curved S-funnel followed by a conventional ion funnel, improves ion transmission efficiency while reducing chemical noise through selective ion focusing. The performance of the dual ion funnel interface was systematically compared to standard interface on a TSQ Vantage platform across samples with different levels of complexity. The dual funnel interface demonstrated to provide up to 25-fold improvement in sensitivity across a wide range of protein concentrations in different biological matrices (low complex mouse macrophage and high complex human cells). Critically, enhanced sensitivity was accompanied by increased analytical reproducibility with lower coefficient of variations. Most importantly, the dual funnel interface enabled reliable quantification of low-abundance proteins that were barely detectable or not detected by the standard interface, extending analysis to single-cell equivalent amounts while maintaining excellent reproducibility. These results demonstrate that the dual funnel interface addresses the critical bottleneck in quantitative targeted proteomics, providing a technological foundation for ultrasensitive targeted SCP that requires both high sensitivity and robust quantitative performance.

## INTRODUCTION

Single-cell proteomics (SCP) has gained attention as a transformative approach to understanding protein expression patterns in individual cells, providing insights into cellular heterogeneity and disease biology.^1, 2^ Unlike traditional bulk proteomics approaches that analyze protein expression from 1,000s of cells, SCP reveals cell-to-cell variability and rare cell populations that are often masked in traditional bulk proteomics.^2^ This capability is particularly crucial for understanding complex biological systems such as immune responses, tumor microenvironments, and developmental transitions, where cellular diversity drives functional outcomes.^3, 4^

A major challenge inherent to SCP primarily stems from the limited protein content of individual cells, typically at the picogram levels.^5^ Mass spectrometry (MS)-based approaches have emerged as a leading technology for SCP due to their ability to provide high molecular specificity, sensitivity, and quantitative capabilities.^6^ Recent advances in MS have enabled researchers to minimize sample losses by improving sample handling and achieving improved data completeness.^7^ However, the detection and quantification of a majority of proteins from single cells remains challenging for current analytical capabilities, driving the need for continuous innovations in both instrumentation and methodology.^7^

Selected reaction monitoring (SRM) using easily accessible triple quadrupole (QqQ) MS is highly effective for targeted SCP studies due to several inherent advantages. SRM operates through targeted monitoring of specific peptide transitions with high precision and selectivity, enabling quantification of low-abundance proteins even in highly complex biological matrices with minimal sample amounts.^8–11^ The exceptional reproducibility and robustness of SRM make it ideally suited for applications requiring consistent quantification across numerous individual cells or limited clinical samples.^8, 9, 12^ Unlike discovery-based methods such as data-dependent acquisition (DDA) or data-independent acquisition (DIA), which prioritize proteome coverage, SRM focuses on predefined targets with exceptional sensitivity and precision.^12, 13^ This targeted approach is especially valuable for studying specific signaling pathways, validating biomarker candidates, or monitoring therapeutic responses, where reliable quantification of key proteins is more important than comprehensive proteome coverage.^12, 13^ Additionally, SRM multiplexing capabilities enable simultaneous monitoring of multiple peptide transitions, facilitating efficient analysis of protein networks or pathway components in a single experiment.^12^

Despite its many advantages, the performance of SRM and other MS-based approaches in SCP remains fundamentally constrained by the MS sensitivity limitations.^14^ Current MS platforms typically require femtomole to attomole levels of protein for reliable detection and quantification, yet low-abundance proteins in single cells may be present at only fewer than 1,000 copies per cell.^15^ The detection and quantification of proteins from single cells is inherently challenging due to at least one order of magnitude short for MS sensitivity. Additionally, signal losses during ionization and transmission through the MS further compound these challenges, requiring the need for continuous innovations in MS instrumentation, particularly in improving ion transmission efficiency and detection sensitivity.^16^ Enhancing sensitivity to reliably detect such ultra-low protein amounts remains a critical focus in advancing SCP.

Recent advances in MS instrumentation have addressed some of these limitations, with innovations such as ion transmission devices and interface designs to improve ion transfer from subambient pressure regions to the vacuum required for MS.^16^ The electrodynamic ion funnel interface design has proven to be effective for capturing and focusing ions over a wide pressure range (1-30 Torr) and has been shown to provide over an order of magnitude enhancement in sensitivity compared to the standard capillary configurations.^17–19^ The evolution from single-stage to dual-stage ion funnel systems, combined with multicapillary inlets, has further enhanced ion transmission efficiency, resulting in improved signal intensity for SRM applications.^16, 20^ However, it has never been demonstrated to integrate the dual-stage ion funnel interface (i.e., dual funnel interface) with conventional QqQ MS for ultrasensitive targeted SCP as well as targeted proteomics analysis of low-input samples in a cost-effective, easily accessible manner.

In this study, we report on the development and comprehensive evaluation of a dual ion funnel interface designed to address the critical sensitivity limitations that constrain current SCP applications. The interface combines a curved S-funnel design with a conventional ion funnel in a tandem configuration specifically engineered to reduce the source contamination and interferences from ESI droplet residue and other poorly focused neutrals when coupled with an in-line injection capillary. We conducted systematic performance evaluations of the dual funnel interface by coupling it to QqQ MS using three experimental systems with different complexity and biological relevance. Our results demonstrate that the dual ion funnel interface achieves improvements in sensitivity (up to 25-fold), quantitative accuracy, and analytical reproducibility relative to a standard S-lens interface while maintaining full compatibility with existing instrumentation and analytical workflows. These advances address the critical technological barrier in sensitivity for targeted proteomics and establish a foundation for routine, ultrasensitive, quantitative targeted SCP.

## EXPERIMENTAL SECTION

### Chemicals and Materials

All chemicals were purchased from Sigma-Aldrich (St. Louis, MO) and Thermo Fisher (Waltham, MA) unless otherwise stated. For initial tuning of the instrument, the triple quadrupole calibration stock solution (25 µM polytyrosine-1,3,6 in 1:1 water/methanol with 0.1% formic acid, Thermo Scientific Pierce, Rockford, IL) was used to tune the standard Thermo source. The same calibration solution was also used to optimize the RF amplitude and DC gradient utilized on the dual ion funnel HPIF/IF interface.

### Sample Preparation

Mouse macrophage cell digests (RAW 264.7) were cultured, lysed, and digested with trypsin following standard proteomics protocols. The resulting digests were serially diluted to generate five different concentrations for sensitivity assessment experiments. For HeLa cell digest preparation, Pierce HeLa Protein Digest Standard (Thermo Fisher) was reconstituted according to manufacturer instructions and diluted to prepare eight different concentrations (0, 0.1, 0.5, 1.0, 2.5, 5.0, 12.5 and 25.0 ng/µL) for LOD/LOQ determination experiments. Each concentration was spiked with synthetic crude heavy isotope-labeled EGFR pathway peptides at a concentration of 25 fmol/µL for each peptide.

For SILAC-labeled HeLa cell preparation, HeLa cells (ATCC; CRM-CCL-2) were cultured in DMEM for SILAC (Thermo Fisher) supplemented with 10% fetal bovine serum (dialyzed, One Shot™ format, Thermo Fisher), 2 mM L-glutamine, and 1% penicillin-streptomycin. For heavy labeling, cells were grown in medium supplemented with L-Arginine-HCl (^13^C_6_, ^15^N_4_) and L-Lysine-2HCl (^13^C_6_, ^15^N_2_) at 0.1 mg/mL. Cells were cultured for at least 40 doublings to ensure complete isotope incorporation before harvesting. Prior to harvesting, cells were rinsed twice with ice-cold PBS, and cell pellets were stored at –80°C. For sample preparation, cell pellets were lysed in 100 mM ammonium bicarbonate (pH 7.8) containing 8 M urea and 75 mM NaCl using repeated vortexing and bath sonication. After centrifugation (14,000 × *g*, 10 min, RT), protein concentration was determined by BCA assay (Thermo Fisher). Samples were reduced with 5 mM DTT (1 h, 37 °C, 800 rpm, Thermo Fisher), alkylated with 10 mM iodoacetamide (1 h, RT, dark, 800 rpm, Thermo Fisher), and diluted 8-fold with 50 mM ammonium bicarbonate before overnight trypsin digestion (1:50 ratio, 37 °C, 800 rpm). Digested samples were acidified with 0.1% TFA and desalted using 100 mg Discovery C18 SPE columns (Millipore Sigma, St. Louis, MO) following manufacturer protocols. Final peptide concentrations were determined by BCA assay after SpeedVac drying and resuspension in 0.1% FA.

### Dual Ion Funnel Interface

This study was performed on a TSQ Vantage triple-stage quadrupole mass spectrometer (Thermo Scientific, Waltham, MA), where the standard S-lens portion of the instrument was replaced with a custom dual-stage ion funnel interface (**Figure 1**). The dual-stage ion funnel configuration consisted of a large bore stainless steel capillary (1 mm i.d., part number U-139, IDEX Health & Science, LLC), an S-shaped high-pressure ion funnel (HPIF, 9.5 Torr, 1.05 MHz, 220 Vp-p),^21^ and a second conventional ion funnel (IF, 1 Torr, 940 kHz, 120 Vp-p).^22^ Conductance-limiting orifices (2.5 mm i.d.) were placed after each ion funnel. The S-funnel was comprised of 169 ring electrodes (0.5 mm thickness, 0.5 mm spacing, 169 mm length) and possessed entrance and exit orifice i.d. of 20.0 mm and 2.5 mm, respectively. The S-funnel possessed four bends (forming an ‘S’ shape) that were designed to decrease the transmission of neutral particles and charged droplets by eliminating line-of-sight to the next pressure chamber. The second funnel is a conventional one of a straight path. The inner diameters of the entrance and exit electrodes were 25.4 mm and 3.0 mm, respectively. The electrodes and spacers for both ion funnels were made from printed circuit board material (FR4, fiberglass-enforced epoxy). Electrical connections between electrodes were made using conductive, spring-loaded pins (Digikey, Thief River Falls, MN, USA) instead of wires. 180° phase-shifted RF waveforms were applied to adjacent funnel electrodes to provide radial ion confinement, and DC gradients were superimposed over the RF to guide ions axially through the funnels. The triple quadrupole calibration stock solution was used to optimize the standard Thermo source and the dual funnel HPIF/IF interface.

**Figure 1.**
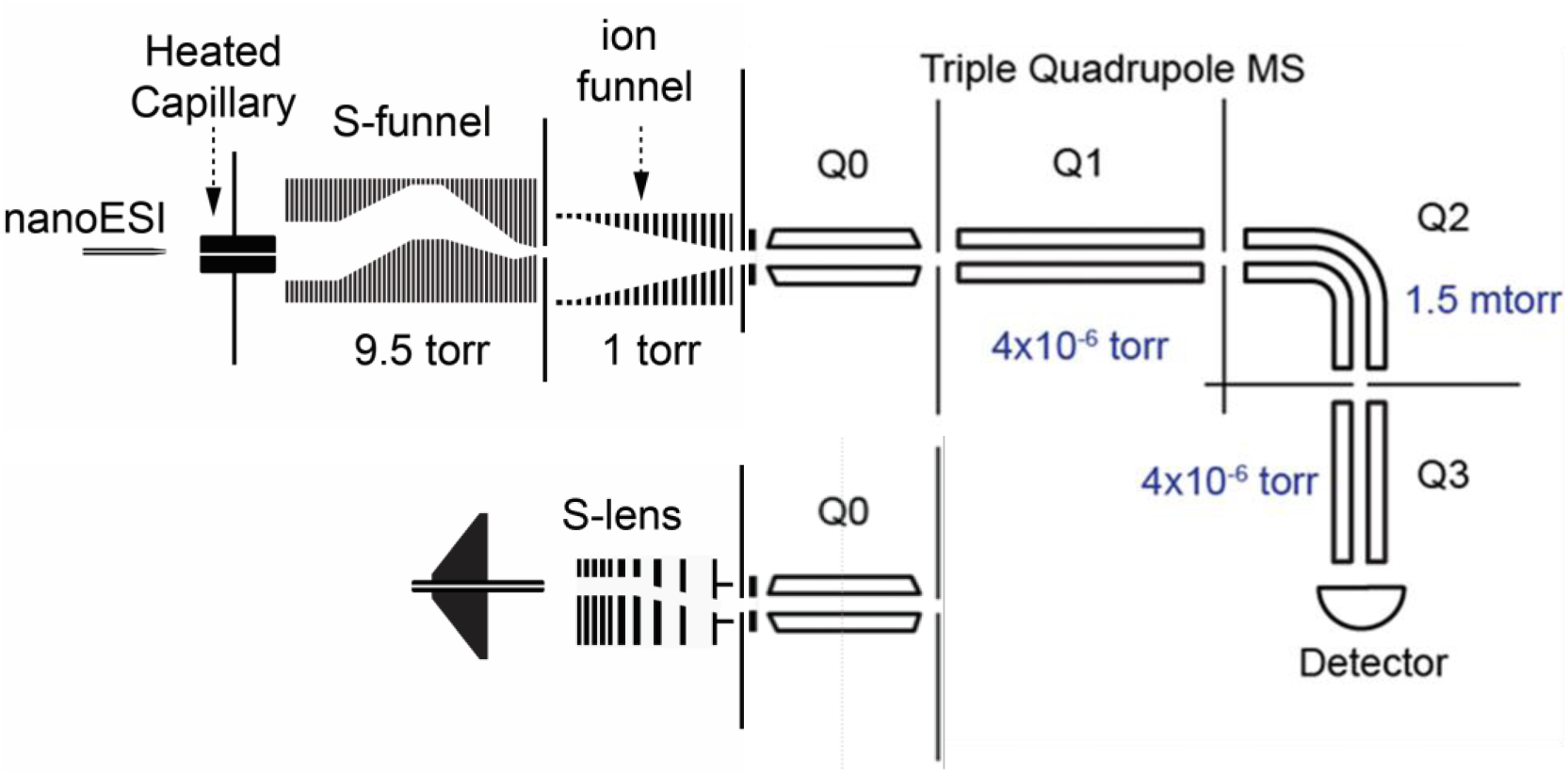
Schematic diagram of the triple quadrupole mass spectrometer platform equipped with the dual ion funnel (HPIF/IF: top) compared to the standard interface (S-lens: bottom).

### LC-SRM-MS Analysis

Three sets of targeted SRM experiments were performed with distinct experimental designs and transition lists. For initial interface performance comparison, RAW 264.7 mouse macrophage digests were analyzed with 236 transitions from 58 unique precursors.

For comprehensive sensitivity and reproducibility assessment, the selection of surrogate peptides for epidermal growth factor receptor (EGFR) pathway proteins and the SRM assays were described previously.^10, 23, 24^ Crude heavy isotope-labeled EGFR pathway peptide standards were spiked into Pierce HeLa Protein Digest Standard (Thermo Scientific, Waltham, MA) at 25 fmol for each peptide. 313 transitions from 50 precursors were monitored and represent 29 EGFR pathway proteins. For SILAC-labeled peptides, heavy isotope-labeled peptide digest from cultured cell lysate (1 ng) was spiked into the unlabeled peptide sample. 456 transitions from 76 unique precursors were monitored and represent 56 proteins. Samples were analyzed using a nanoACQUITY UPLC (Waters Corporation, Milford, MA) coupled to a TSQ Vantage triple-stage quadrupole mass spectrometer. The UPLC M-Class Peptide BEH 130Å, 1.7 μm C18 column (100 μm i.d. × 10 cm) was connected to a hydrofluoric acid-etched 20 μm i.d. fused-silica electrospray emitter^25^ via a stainless metal union. Solvents used were 0.1% formic acid in water (mobile phase A) and 0.1% formic acid in 90% acetonitrile (mobile phase B). 1 μL peptide sample was directly loaded onto the BEH C18 column from a vial without using a trapping column. Sample loading and separation were performed at flow rates of 350 and 300 nL/min, respectively. The binary LC gradient was used: 5–20% B in 26 min, 20–25% B in 10 min, 25–40% B in 8 min, 40–95% B in 1 min and at 95% B for 7 min for a total of 52 min. The analytical column was re-equilibrated at 99.5% A for 8 min. The TSQ Vantage mass spectrometer was operated with ion spray voltages of 2400 ± 100 V, a capillary offset voltage of 35 V, a skimmer offset voltage of −4 V, and a capillary inlet temperature of 340 °C. The tube lens voltages were obtained from automatic tuning and calibration without further optimization. The retention time scheduled SRM mode was applied for SRM data collection with the scan window of ≥ 6 min. The cycle time was set to 1 s, and the dwell time for each transition was automatically adjusted depending on the number of transitions scanned at different retention time windows. A minimal dwell time 10 ms was used for each SRM transition. All the EGFR pathway proteins were simultaneously monitored in a single LC-SRM analysis.

### LC-SRM Data Analysis

SRM data acquired on the TSQ Vantage were analyzed using Skyline software. Matrix inferences from co-eluting peptides with a transition that fell within the mass width of Q1 and Q3 were detected by deviation from the expected L/H SRM signal ratios. The best transition for each peptide was used for quantification, and the L/H SRM ratios were used to generate calibration curves. Peak detection and integration were based on retention time alignment and relative peak intensity ratios across multiple transitions between light and heavy peptides. All data were manually inspected to ensure correct peak detection and accurate integration.

Signal-to-noise (S/N) ratios were calculated by dividing the intensity of the peak apex by the highest background noise in a retention time window of ±15 s for the target peptides. Limit of detection (LOD) and limit of quantification (LOQ) were defined as the lowest concentration point of each target peptide at which the S/N ratio was at least 3 and 10, respectively. For stringent determination of LOQ, two additional criteria were implemented: 1) Peptide response falls within the linear dynamic range; 2) CV at LOQ is below 20%. The RAW data from TSQ Vantage were loaded into Skyline software to display graphs of extracted ion chromatograms (XICs) of multiple transitions of target proteins monitored.

### Statistics and Reproducibility

Standard deviation (SD) and coefficient of variation (CV) were calculated from up to four technical replicates. Statistical analysis of the data was carried out using the R software environment (version 4.3.3) with additional software packages.

## RESULTS AND DISCUSSION

To address the ion transmission limitations of conventional interfaces, we designed a dual ion funnel interface device. This device, composed of tandem S-funnel and standard funnel, was engineered to enhance ion transmission for improved detection sensitivity compared to a standard interface when coupled to the TSQ Vantage triple quadrupole mass spectrometer (**Figure 1**). The first funnel that ions pass through after ionization is the S-funnel, which features a curved geometry with a non-linear ion path that allows only ions to follow the curved center axis, while non-ionic species are efficiently removed between the electrodes through cutouts in the spacers. This first-stage high-pressure ion funnel (HPIF) operates at approximately 9.5 Torr. The second conventional ion funnel (IF), positioned immediately behind the HPIF, maintains a pressure of 1 Torr, ensuring high ion transmission efficiency through the interface. To comprehensively evaluate the performance improvement of the dual ion funnel interface (**Figure 1** *top*) for targeted SCP, we conducted systematic comparisons with the standard interface (**Figure 1** *bottom*) across multiple analytical metrics including sensitivity, detection limits, and quantitative precision.

### Initial Sensitivity Assessment Using Low-Complexity Matrix

Initial performance assessment focused on establishing the sensitivity improvement using mouse macrophage cell digests as a low-complexity matrix model. This cell line was selected as an initial test system due to its well-characterized proteome and low complexity, allowing for reliable assessment of fundamental performance differences between the interfaces. Five serially diluted mouse macrophage cell lysate digests (RAW 264.7 cells) were analyzed, and a systematic side-by-side comparison between the dual funnel interface and the standard interface was performed under their optimal conditions. A total of 236 transitions from 58 unique precursors (3-5 fragment ions per precursor) were monitored in SRM mode to cover a wide dynamic digest input amounts. The dual funnel interface demonstrated substantial sensitivity improvements across all monitored transitions. On average, peak areas were enhanced approximately 33-fold with the dual funnel interface, with individual peptide improvements ranging from 15- to 83-fold. **Figure 2** illustrates the performance analysis of a representative peptide that exhibits the average fold-change improvement observed across all the different digest input amounts. The extracted ion chromatograms (XICs) demonstrate the dramatic signal enhancement achieved with the dual funnel interface (**Figure 2A**), while both interfaces maintained excellent linearity (R^2^ > 0.98) in their calibration curves (**Figure 2B**).

**Figure 2.**
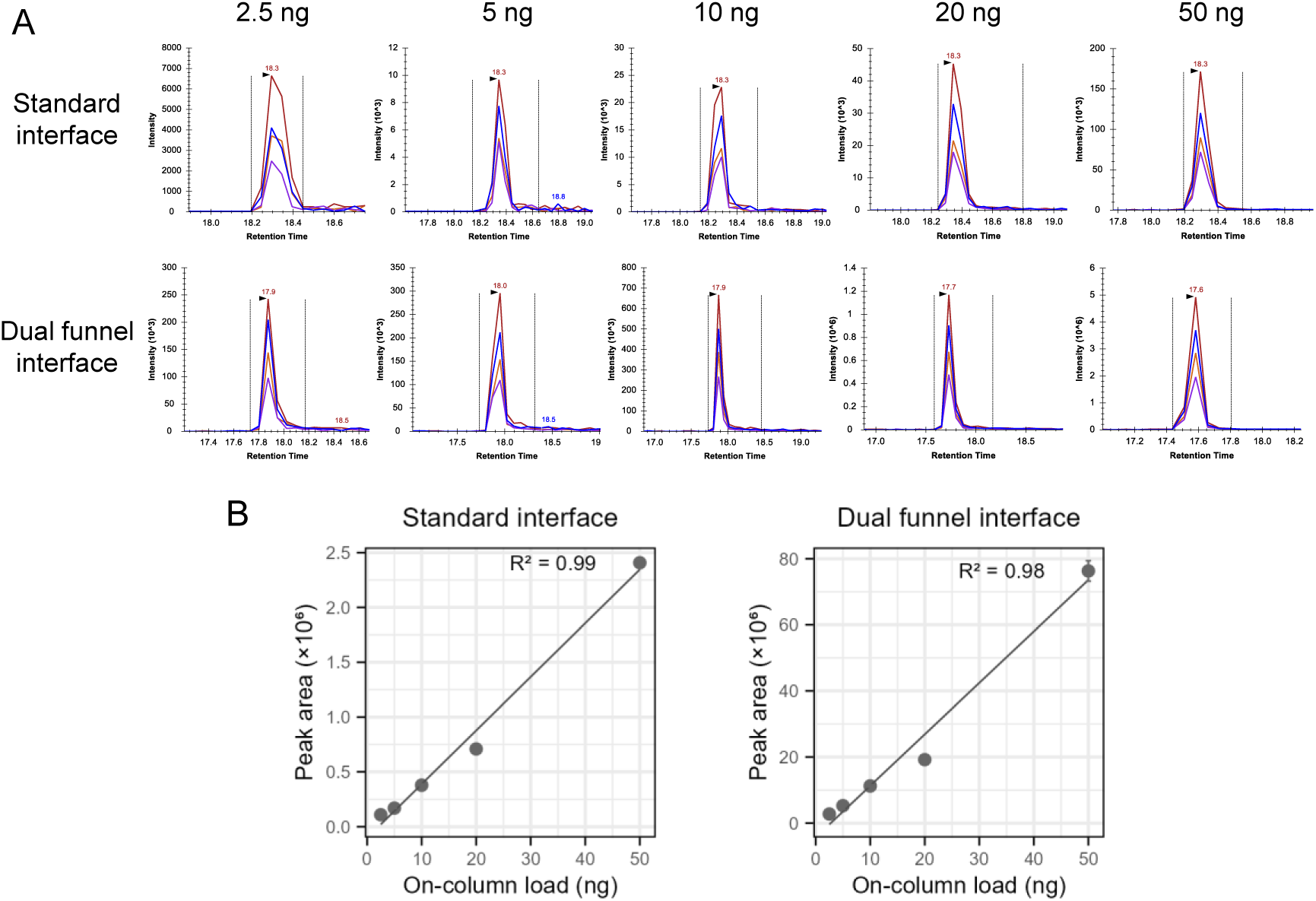
Sensitivity improvement of dual funnel interface in low-complexity matrix. Representative peptide GYSFTTTAER (derived from actin) demonstrating the average ∼33-fold sensitivity improvement observed across all monitored peptides from mouse macrophage cell (RAW 264.7) digests. **(A)** Extracted ion chromatograms (XICs) of transitions comparing standard interface (top) and dual funnel interface (bottom) across all tested digest concentrations. **(B)** Calibration curves showing peak area versus sample concentration for both interfaces. Both interfaces demonstrate excellent linearity (R^2^ > 0.98), with the dual funnel interface providing consistently enhanced signal intensity. Error bars represent standard deviations of the measurements.

Analysis of the fold-change distribution across all monitored peptides revealed significant improvement in the sensitivity for the dual funnel interface. Peptides with relatively higher signal intensities showed moderate fold-change improvements, while lower abundance peptides demonstrated substantial enhancements. While some peptides, such as the representative example in **Figure 2**, maintained consistent fold-change across digest input amounts, other peptides showed concentration-dependent variation in fold-change improvements, likely reflecting the complex interplay between individual peptide properties and complex matrix effects. This differential improvement pattern indicates that the enhanced sensitivity of the dual funnel interface results in an expanded dynamic range, effectively bringing previously low-abundance peptides into reliably detectable ranges. This finding suggests that the dual funnel interface would be particularly beneficial for sensitive quantification of low-abundance proteins with significant improvement in detection sensitivity for low-complex samples.

### Quantification Performance Assessment Using Complex Biological Matrix with Spiked-in Heavy Isotope-labeled Peptide Standards

To provide a more rigorous and biologically relevant evaluation, the sensitivity and precision of the dual funnel interface were further assessed through targeted quantification of EGFR pathway proteins in complex HeLa cell digests. We selected EGFR pathway proteins for this analysis due to several key advantages: the availability of heavy peptide internal standards with established SRM assays and well-characterized protein copy numbers per cell from previous studies.^10, 23, 24^ Eight different concentrations of HeLa digests (0, 0.1, 0.5, 1.0, 2.5, 5.0, 12.5, and 25.0 ng/µL) were spiked with synthetic heavy isotope-labeled EGFR pathway crude peptides (25 fmol/µL) as internal standards. With the spiked-in crude heavy peptide standards at 25 fmol per peptide, the on-column amounts of HeLa digests were 0, 0.1, 0.5, 1.0, 2.5, 5.0, 12.5, and 25.0 ng, which are equivalent to 0, 0.4, 2, 4, 10, 20, 50, and 100 cells, respectively. A total of 313 transitions from 50 unique precursors (29 EGFR pathway proteins) were monitored in SRM mode to cover a wide dynamic protein abundance range. This design enabled comprehensive evaluation of detection sensitivity through LOD and LOQ determinations, while technical reproducibility was assessed by calculating the coefficient of variation (CV) of the light-to-heavy (L/H) peptide peak area ratios across replicates. Comparative performance evaluation was performed on pathway proteins representing different abundance ranges (**Table 1**): high-abundance proteins (>500,000 copies/cell: PEBP1, GRB2), moderate-abundance proteins (100,000-500,000 copies/cell: MAP2K1, NRAS, PTPN11, SHC1), and low-abundance proteins (<100,000 copies/cell: ARAF, RASA1, ADAM17).^24^

**Table 1.**
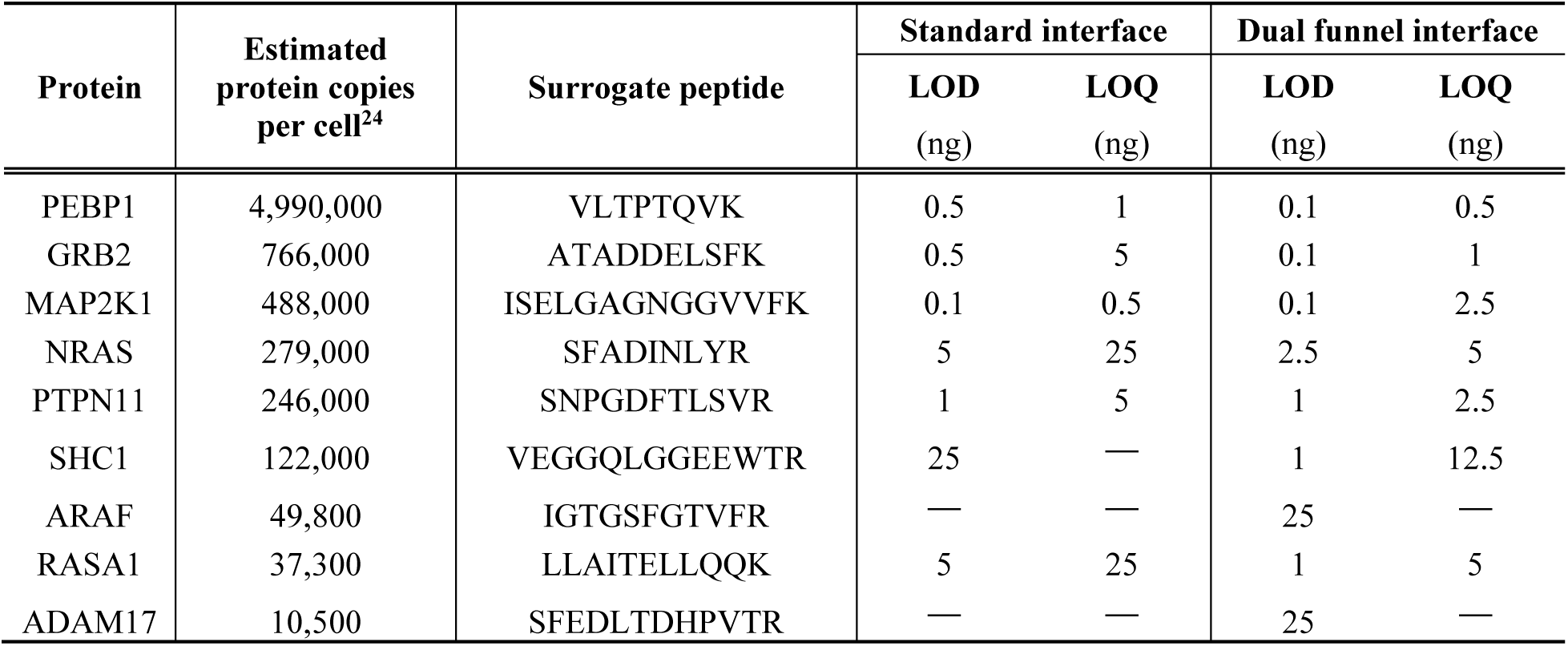
Detection and quantification limits of selected EGFR pathway peptides. Limits of detection (LOD) and limits of quantification (LOQ) for representative peptides from different protein abundance categories (high, moderate, and low) in HeLa cell digests. Comparison between the standard interface and dual funnel interface demonstrates systematic improvements in detection sensitivity across all abundance ranges.

### Detection and Quantification Performance Assessment

**Figure 3** depicts representative XICs and calibration curves for peptides from proteins across this abundance spectrum, demonstrating the systematic performance improvements achieved by the dual funnel interface. For a high-abundance target, despite the readily detectable nature of high-abundance targets, the dual funnel interface still provided substantial sensitivity enhancement. Both interfaces successfully detected VLTPTQVK (derived from PEBP1, ∼4,990,000 copies/cell), but the dual funnel interface provided superior performance. At 0.1 ng HeLa digests, the dual funnel interface achieved S/N of 4 compared to 2 for the standard interface, while enabling LOD improvement from 0.5 to 0.1 ng and LOQ improvement from 1 to 0.5 ng—representing 5-fold and 2-fold improvements, respectively (**Figure 3A** and **Table 1**). Similarly, ATADDELSFK (derived from GRB2, ∼766,000 copies/cell) showed LOD improvement from 0.5 to 0.1 ng and LOQ improvement from 5 to 1 ng (5-fold improvement for both metrics) (**Table 1**). The performance improvements are also observed for moderate-abundance proteins.

**Figure 3.**
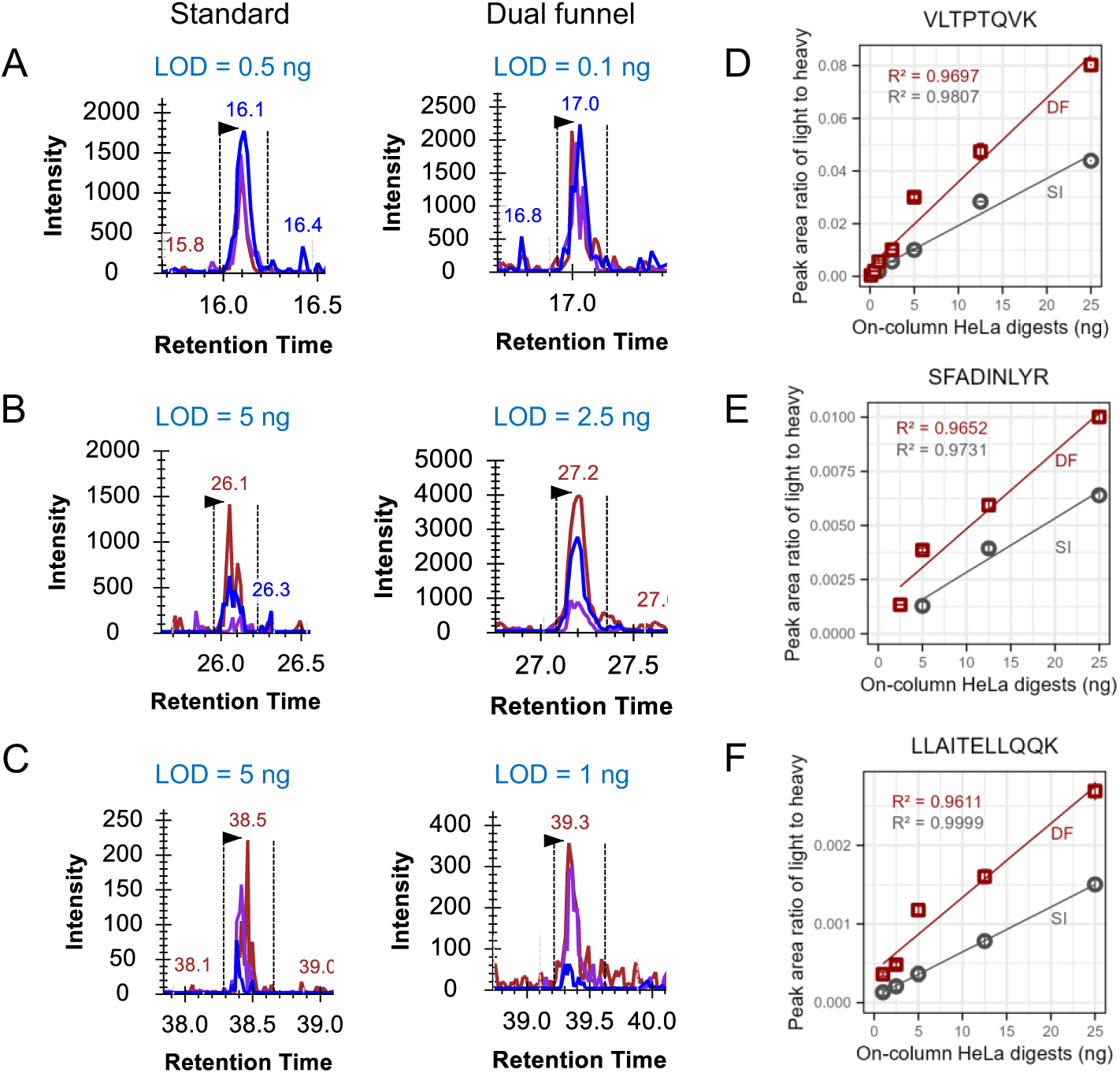
Detection and quantification performance across protein abundance ranges in complex biological matrix. Representative peptides from EGFR pathway proteins demonstrated the enhanced sensitivity and quantification accuracy of the dual funnel interface in HeLa cell digests spiked with heavy isotope-labeled internal standards (25 fmol). **(A-C)** XICs of light peptide transitions at each LOD comparing standard interface (left) and dual funnel interface (right). **(A)** VLTPTQVK from high-abundance PEBP1 (4,990,000 copies/cell). **(B)** SFADINLYR from moderate-abundance NRAS (279,000 copies/cell). **(C)** LLAITELLQQK from low-abundance RASA1 (∼37,300 copies/cell). **(D-F)** Calibration curves showing light-to-heavy peak area ratios versus HeLa digest concentration for the corresponding peptides in panels **A-C**, respectively. The dual funnel interface demonstrates superior sensitivity across all abundance ranges and enhanced quantification accuracy with closer agreement to expected theoretical ratios. Error bars represent standard deviations. SI: standard interface; DF: dual funnel interface.

SFADINLYR (derived from NRAS, ∼279,000 copies/cell) achieved LOD improvement from 5 to 2.5 ng and LOQ improvement from 25 to 5 ng—representing 2-fold and 5-fold improvements, respectively (**Figure 3B** and **Table 1**). Most significantly, for low-abundance proteins, the dual funnel interface enabled detection of targets that were below the detection capabilities of the standard interface (**Table 1**). LLAITELLQQK (derived from RASA1, ∼37,300 copies/cell) showed LOD improvement from 5 to 1 ng and LOQ improvement from 25 to 5 ng (5-fold improvement for both metrics) (**Figure 3C** and **Table 1**). IGTGSFGTVFR (derived from ARAF, ∼49,800 copies/cell) and SFEDLTDHPVTR (derived from ADAM17, ∼10,500 copies/cell) remained below detection limits for the standard interface (S/N = 2 at 25 ng) while achieving LOD of 25 ng with the dual funnel interface (S/N = 4).

Quantification performance analysis revealed that the dual funnel interface substantially improved LOD and LOQ values directly translated into enhanced quantitative capabilities with broader dynamic ranges. The dual funnel interface showed reliable quantification accuracy by precisely reflecting the expected proportional changes in L/H area ratios corresponding to light peptide concentration differences (**Figure 3D-F**). The dual funnel interface produced the L/H area ratios that closely matched these fold-changes of the light peptide inputs with excellent quantitative linearity (R^2^ > 0.95). In contrast, the limited quantification range of the standard interface, with LOQ values often at or near the maximum of tested peptide inputs, severely restricted the ability to assess quantitative accuracy and concentration-dependent responses. The enhanced quantification performance of the dual funnel interface is particularly critical for quantitative targeted proteomics applications, where precise determination of protein abundance differences between samples depends on the linear relationship between actual protein concentrations and the measured L/H area ratios.

### Analytical Reproducibility at Detection Limits

To assess the analytical reliability at the critical performance thresholds, we evaluated reproducibility at LOD levels, which is particularly important for low-abundance protein quantification, where measurements often operate near detection limits. Analysis of CVs of the L/H area ratios at respective LOD levels revealed excellent reproducibility for the dual funnel interface across most protein abundance categories. For proteins where both interfaces achieved measurable LODs, the dual funnel interface consistently enabled detection at much lower concentrations while maintaining high reproducibility. For low-abundance proteins, the dual funnel interface achieved acceptable reproducibility with CVs below 15% at LOD (IGTGSFGTVFR: 2.3%, LLAITELLQQK: 9.3%, SFEDLTDHPVTR: 13.5%). This represents a significant expansion of the quantitation dynamic range, extending reliable detection to the low-abundance protein range that constitutes a significant portion of the cellular proteome.

To demonstrate the practical advantage of improved detection limits, we compared the CVs from both interfaces at digest input amounts corresponding to the standard interface LODs. At these digest inputs, the dual funnel interface operates well above its own detection limits, resulting in significantly improved reproducibility compared to the standard interface. For the standard interface, the individual peptide CVs at their LODs ranged from 7.2% to 83.8% with an average of 43.6%, whereas the dual funnel interface achieved the CVs of 1.0% to 35.8% with an average of 15.4% at these same digest inputs. These results demonstrate that the enhanced detection sensitivity provided by the dual funnel interface translates into more reliable quantification in the digest input range where the standard interface struggles to provide consistent results. When measured at the LODs of the dual funnel interface, the CVs ranged from 2.3% to 35.8% with an average of 14.0%, demonstrating maintained reproducibility even at substantially lower peptide amounts.

### Comprehensive Analytical Reproducibility Assessment

Beyond the reproducibility at detection limits, we evaluated analytical reproducibility across all experimental conditions using two complementary approaches to establish the overall analytical reliability of each interface. First, the technical system reproducibility of both interfaces was evaluated using all 50 monitored crude heavy peptide internal standards spiked at constant concentrations across all experimental conditions. Since heavy peptides were maintained at identical concentrations throughout all measurements, they served as direct indicators of system-level reproducibility. Both the standard and dual funnel interfaces demonstrated similar technical reproducibility with a mean heavy peptide CV of 11.3% for the standard interface and 11.1% for the dual funnel (**Figure 4A**).

**Figure 4.**
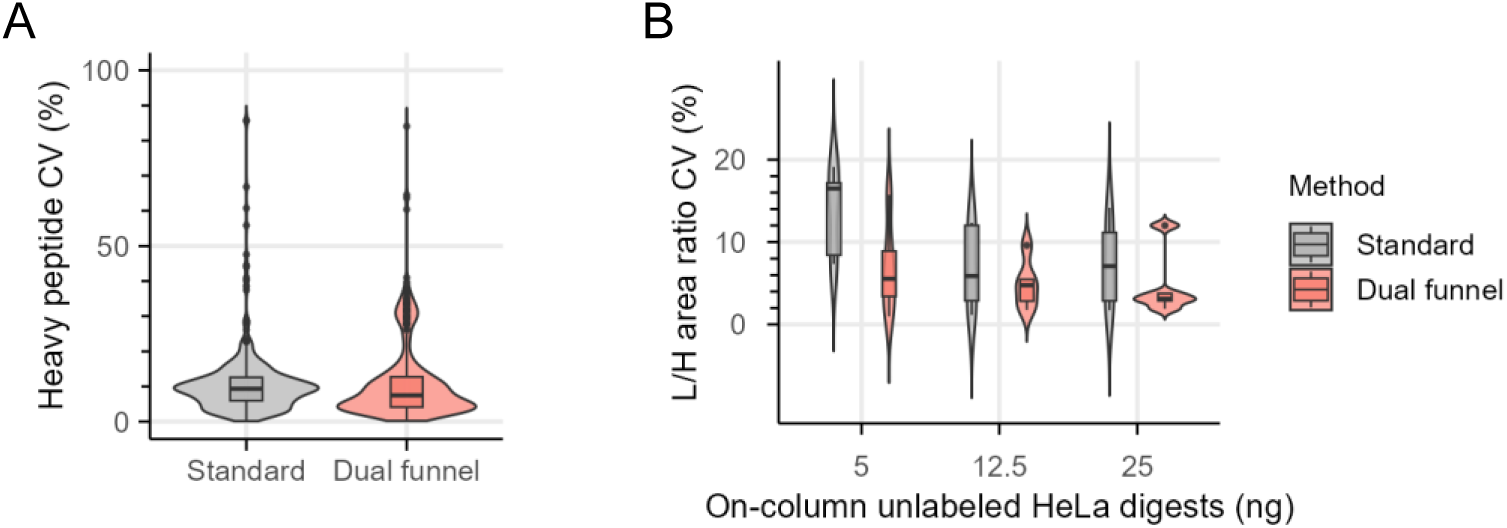
Analytical reproducibility comparison between standard and dual funnel interfaces. **(A)** Violin plot showing the distribution of heavy peptide peak area coefficient of variation (CV) values for the standard interface and dual funnel interface. Heavy peptides were spiked at constant amounts (25 fmol) across all experimental conditions to assess technical system reproducibility. Each violin represents the distribution of CV values across all 50 monitored heavy peptides. **(B)** Violin plot comparing light-to-heavy (L/H) area ratio CV distributions between the two interfaces across HeLa digest inputs from 5 to 25 ng. Only peptide-concentration combinations quantifiable by both interfaces (above their respective LOQs) were compared (5 peptides per concentration). The dual funnel interface showed consistently lower median CVs and tighter distributions. Box plots within violins indicate median, quartiles, and outliers.

Second, a comprehensive analysis of L/H area ratio CVs was performed using only peptides that were quantifiable in both interfaces across matched experimental conditions, ensuring a fair comparison between the two interfaces. Specifically, for each peptide-concentration combination, measurements were included only when the digest input amount exceeded the respective LOQ of each interface (i.e., standard interface measurements required concentration ≥ LOQ of standard, while dual funnel interface measurements required concentration ≥ LOQ of dual funnel). This conservative approach excluded measurements below the detection threshold of each interface and only included peptide-concentration combinations where both interfaces independently achieved reliable quantification above their respective performance limits. This matched-peptide analysis revealed substantial and consistent improvements in quantitative reproducibility with the dual funnel interface across all digest input amounts, with particularly pronounced improvements at low-to-intermediate digest inputs where quantification approached the LOQ limits (**Figure 4B**).

These comprehensive assessments demonstrate that the dual funnel interface provides systematic improvements across all evaluated metrics: detection sensitivity, dynamic range, and analytical reproducibility. The unique strength of the dual funnel interface lies in significantly expanding the analytical capabilities for low-abundance protein analysis while maintaining high reproducibility. The dual funnel interface achieved 2- to 25-fold improvements in detection limits across all tested peptides and critically extended reliable quantification to medium-to-low abundance proteins that were beyond the quantitative range of the standard interface. Even for ultra-low abundance proteins where quantification remained challenging, the dual funnel interface enabled consistent detection at biologically relevant concentrations. This combination of enhanced sensitivity and robust reproducibility positions the dual funnel interface as a transformative platform for quantitative targeted SCP applications.

### Ultrasensitive Targeted Quantification Using Dual Funnel Interface for Broad Applications

Having demonstrated improved performance of the dual funnel interface using synthetic heavy peptide internal standards for targeted EGFR pathway analysis, we next sought to establish a more versatile and cost-effective approach for single-cell level targeted proteomics. We employed SILAC-labeled cell lysate digests as a universal internal standard (SILAC-UIS), which offers significant advantages over conventional synthetic heavy peptide standards: it enables multiplexed quantification without requiring individual standards for each target protein, serves as an effective carrier to minimize sample loss in low-input analyses similar to our previously reported carrier-assisted approach, and substantially reduces cost.^10, 11^

### Optimization of Universal Internal Standard Implementation for Low-Input Sample Analysis

To determine the optimal SILAC-UIS implementation strategy, we first performed proteome-wide evaluation using data-independent acquisition (DIA) mode on an Orbitrap Exploris 480 MS platform. We tested unlabeled HeLa cell digests ranging from 0.25 to 10 ng (equivalent to 1-40 cells in protein content) spiked with 1 ng of SILAC-labeled HeLa digest. Representative XICs for VLTPTQVK (derived from PEBP1) demonstrated quantifiable signal intensities and excellent linearity across the tested range. As expected, comprehensive DIA characterization revealed that the number of identified peptides and protein groups increased progressively with higher lysate amounts. Despite this expected trend, SILAC-UIS maintained quantitative reliability across all tested amounts, as demonstrated by consistent log₂ intensity distributions, stable SILAC ratios, and precise fold-change measurements. Importantly, even at the lowest input (0.25 ng, single-cell equivalent), sufficient peptides and proteins were identified for reliable quantitative analysis. Based on the balance between light peptide signal and the L/H ratios across 1-40 cell equivalent range, we selected 1 ng of SILAC-labeled HeLa digest as the optimal spike-in level for subsequent targeted SRM analysis with the dual funnel interface.

### Single-Cell Level Targeted Proteomics Using Dual Funnel Interface-based QqQ MS Coupled with SILAC-UIS

With the determined spike-in level for SILAC-UIS, we next evaluated the quantitative performance of the integrated dual funnel interface with SILAC-UIS for single-cell level targeted proteomics. We monitored 456 transitions from 76 unique precursors representing 56 proteins spanning a broad range of cellular abundances (54,500 to ∼33 million copies per HeLa cell).^24^ To assess quantitative performance across the low-input range, unlabeled HeLa digests ranging from 0.2 to 50 ng (equivalent to 1-200 cells) were spiked with the optimized 1 ng of SILAC-labeled HeLa digest and analyzed using the dual funnel interface-based QqQ MS platform in SRM mode. The dual funnel interface demonstrated exceptional sensitivity for single-cell level analysis. Representative XICs for three peptides spanning the abundance range showed robust, quantifiable signals even at the single-cell equivalent input (0.2 ng) (**Figure 5A**). Critically, this performance was achieved in the most complex biological matrix tested in this study, where both the sample and internal standard comprise complete cell lysate digests, representing the full proteome complexity and potential matrix effects encountered in authentic single-cell analyses. Quantitative performance metrics for eight representative peptides are summarized in **Table 2**, demonstrating LOD values ranging from 0.2 to 1 ng and LOQ values of 0.2 to 10 ng. This enabled reliable quantification at or near single-cell levels for most monitored proteins. Based on the ∼25-fold LOD improvements demonstrated in the EGFR pathway analysis, this single-cell level quantification capability represents a substantial improvement over standard QqQ MS platforms.

**Figure 5.**
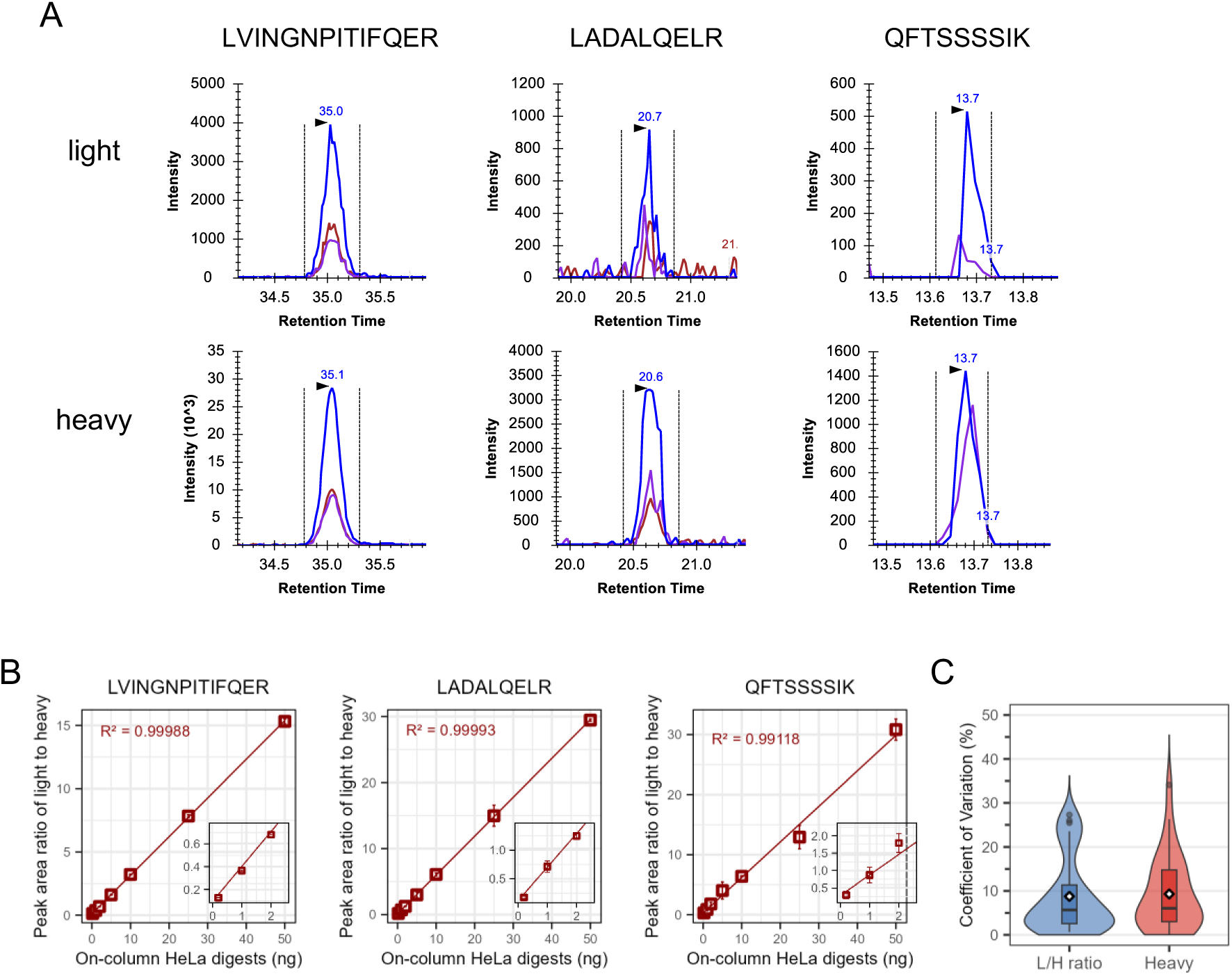
Ultrasensitive peptide quantification using dual funnel interface with SILAC-UIS. **(A)** XICs for LVINGNPITIFQER (derived from GAPDH), LADALQELR (derived from LMNA), and QFTSSSSIK (derived from KRT17) from HeLa cell digests (0.2 ng) spiked with SILAC standards (1 ng) analyzed in SRM mode. **(B)** Calibration curves using the best transitions, with insets showing low-input performance. Error bars represent standard deviations. The dual funnel interface enables reliable quantification with extending to single-cell equivalent amounts. **(C)** CV distributions of L/H ratios and heavy peptide areas for concentrations above LOQ. Box plots indicate median, mean (in white diamond), quartiles, and outliers.

**Table 2.**
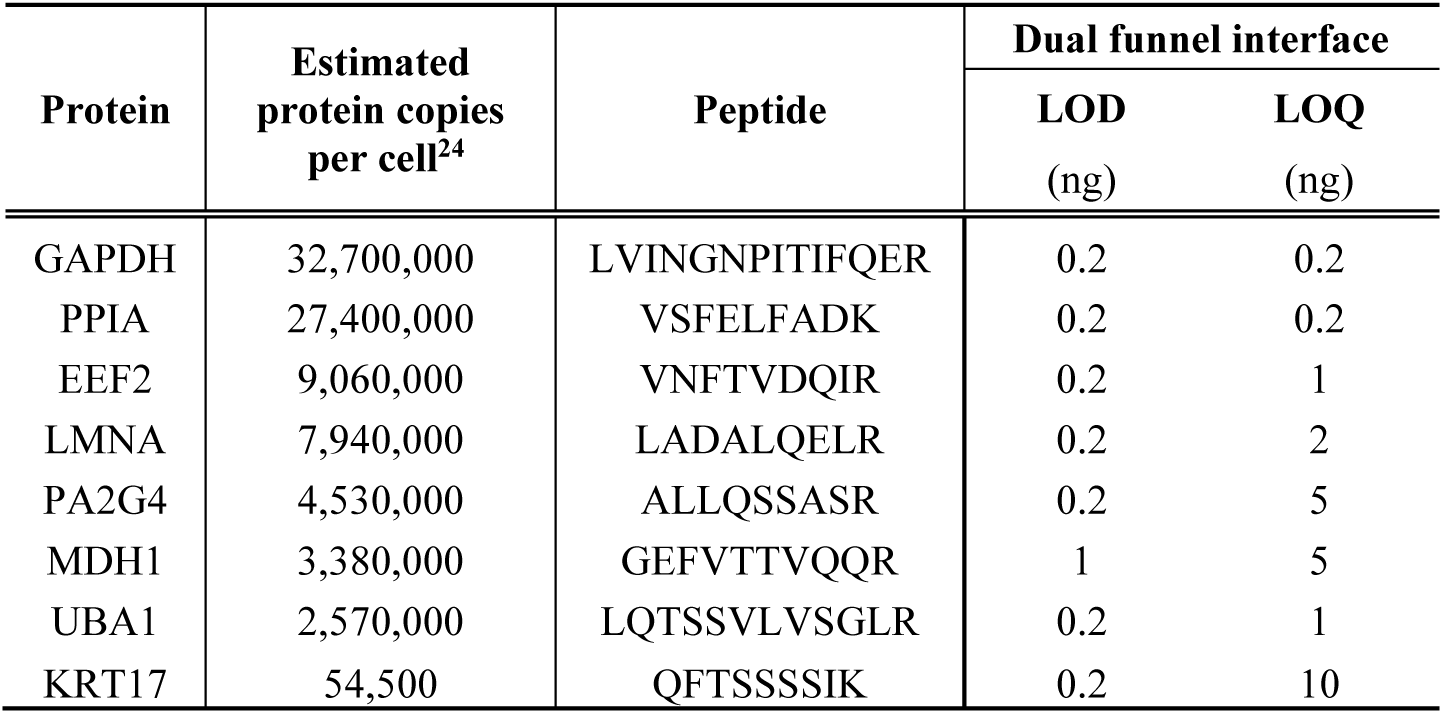
Detection and quantification limits using SILAC-UIS platform with dual funnel interface. LOD and LOQ values for peptides representing proteins covering a wide protein abundance range. Unlabeled HeLa digests (0.2-50 ng) were analyzed with SILAC-labeled HeLa digest (1 ng) as universal internal standards in SRM mode.

Beyond sensitivity, the integrated platform demonstrated excellent quantitative accuracy and precision in this highly complex matrix. Calibration curves constructed from the best-performing transitions for each peptide showed excellent linearity (R² > 0.99) across the tested range, with quantifiable L/H area ratios maintained down to the single-cell level (**Figure 5B**). Notably, the inset plots confirm that accurate concentration-dependent responses are preserved even at ultra-low inputs.

To ensure accurate assessment of quantification performance, reproducibility was evaluated only for peptides achieving quantifiable signals at each input amount. Peptides were considered quantifiable when their best transition met the LOQ criterion (mean S/N ≥ 10 across technical replicates) at the given input level. At the single-cell level (0.2 ng), 31 out of 76 monitored peptides (40%) achieved quantifiable signals and demonstrated excellent L/H area ratio reproducibility with a mean CV of 8.8% (median: 5.7%) (**Figure 5C**). At higher input amounts, over 56 peptides achieved quantifiable signals with mean CVs of 8.0-11.3%, demonstrating robust quantification performance across the low-input range. Heavy peptide signals from the fixed 1 ng SILAC spike-in showed comparable reproducibility (mean CV: 9.3% at 0.2 ng, median: 6.1%) (**Figure 5C**), confirming the robust carrier function of SILAC-UIS in minimizing sample loss across varying sample-to-carrier ratios.

This performance demonstrates that the combination of the dual funnel and SILAC-UIS enables reliable, precise quantification at single-cell equivalent amounts in complex biological matrices. The ability to maintain quantitative accuracy and precision when analyzing 0.2 ng of sample in the presence of 1 ng of SILAC carrier highlights the robustness of this integrated approach under realistic single-cell analysis conditions. Critically, this approach eliminates the need for individual synthetic heavy peptide standards while enabling multiplexed quantification across proteins spanning a wide range of cellular abundances. The demonstrated capability to achieve reproducible quantification at single-cell protein content directly addresses the critical sensitivity bottleneck in targeted SCP and establishes a practical platform for analyzing rare cell populations, early disease states, and limited clinical specimens where sample availability is inherently constrained. Most importantly, this ultrasensitive performance was achieved using standard SRM methodology with the dual funnel interface alone, without requiring specialized sample preparation or signal amplification techniques, making it immediately applicable to existing proteomics workflows.

## CONCLUSIONS

This study highlights the dual ion funnel interface as a novel capability for enhancing the sensitivity of SRM-based targeted proteomics when integrated with conventional QqQ MS. By systematically comparing the dual funnel with standard interfaces for targeted quantification of low-input samples including single-cell level amount, we have demonstrated the superior performance of the dual funnel with significant improvements in sensitivity (up to 25-fold enhancement), quantitative accuracy (closer agreement with theoretical L/H ratios), and analytical reproducibility (∼72% improvement in CVs). Notably, the dual funnel interface enabled reliable detection of low-abundance proteins in low-input samples, including key signaling pathway proteins, thereby overcoming key limitations in current instrumentation. Integration with SILAC-based universal internal standards further validated this capability, achieving detection and quantification at single-cell equivalent amounts (0.2 ng input). The dual funnel interface is fully compatible with conventional QqQ MS and existing targeted proteomics workflows, ensuring rapid adoption by the broad scientific community without substantial infrastructure investment. This advancement establishes the dual funnel interface as a transformative tool for ultrasensitive targeted single-cell proteomics, opening new research frontiers for precise characterization of cellular heterogeneity and improved understanding of single-cell biology.

## ACKNOWLEDGMENTS

This work was supported by NIH R01GM139858 (to T.S.), NIH R33CA287139 (to T.S.), NIH RF1MH128885 (to T.S.) and NIH UH3CA256967 (to T.S.). This project was performed in the Environmental Molecular Sciences Laboratory, a DOE OBER national scientific user facility on the PNNL campus. PNNL is a multiprogram national laboratory operated by Battelle for the DOE under contract DE-AC05-76RL01830. The authors would like to thank Dr. Adam Hollerbach for his helpful comments in reviewing this manuscript.

